# KRT17 stabilizes EPN1 via inhibiting SMURF1-mediated ubiquitination to modulate Wnt/β-catenin signaling output and stem-like traits in ovarian cancer

**DOI:** 10.64898/2026.02.20.707140

**Authors:** Guiqiang Du, Bilan Li, Ruru Zhao, Huan Tong, Yinyan He, Jing Ding

## Abstract

**Background:** Ovarian cancer (OC) progression and chemoresistance are closely linked to dysregulated oncogenic signaling, including Wnt/β-catenin pathways that contribute to cancer stem-like traits. However, the upstream mechanisms connecting cytoskeletal regulation to Wn/β-catenin signaling in OC remain incompletely understood. Keratin 17 (KRT17), a type I intermediate filament protein, has been implicated in tumor progression, but its mechanistic role in OC requires clarification.

**Methods:** Gene Expression Omnibus (GEO) datasets and clinical specimens were analyzed to assess KRT17 and EPN1 expression and prognostic significance. Functional assays, xenograft models, co-immunoprecipitation, ubiquitination analyses, and rescue experiments with wild-type and ubiquitination-resistant EPN1 mutants were performed to investigate molecular mechanisms.

**Results:** KRT17 expression was elevated in OC tissues and correlated with poor patient survival. KRT17 depletion suppressed proliferation, migration, stem-like properties, tumor growth, and cisplatin resistance. Mechanistically, KRT17 interacted with EPN1 and weakened its association with the E3 ligase SMURF1, reducing SMURF1-mediated ubiquitination at lysine 107 and preventing proteasomal degradation. Stabilized EPN1 was associated with increased β-catenin abundance and stemness-associated markers, and enhanced self-renewal capacity.

**Conclusions:** These findings identify a KRT17–EPN1 axis that links intermediate filament dynamics to ubiquitin-dependent regulation of EPN1 stability and Wnt/β-catenin signaling outputs in ovarian cancer.

## 1. Introduction

Cellular signaling networks coordinate crucial cellular functions by receiving input from external signals and processing this information through intracellular control logic. Disruption of both these signaling pathways is a common feature in cancer development, inducing atypical proliferation, migration and transcriptional reprogramming(1). Of these pathways, the Wnt/β-catenin pathway serves as a pivotal signaling center to govern stem-like properties, as β-catenin is typically stabilized and translocated into the nucleus for cell fate determination(2). Continuous activation of the Wnt/β-catenin pathway contributes to increased tumor aggressiveness, which is strongly associated with cancer stemness maintenance(3). One of the most distinctive features of cancer stemness is enhanced self-renewal, cellular plasticity, and capacity for tumor initiation(4). These traits are often supported by signaling-dependent transcriptional responses, as the promotion of stemness related genes (such as SOX2, OCT4, NANOG, and CD133) (5). In ovarian cancer, Wnt/β-catenin signaling promotes malignant progression and has previously been correlated with elevated tumour-initiating potential and therapy resistance(6). However, the upstream pathways linking cytoskeletal reorganization and intracellular adapter proteins to Wnt/β-catenin activation remain poorly characterized.

Keratin 17 (KRT17) belongs to the family of type I intermediate filament protein and has been well-known as a structural component of cytoskeleton(7). Although extensive studies on keratins have shown that they contribute to the shape of cells or tissues, new findings reveal that keratin may play a nonclassical function, such as organizing signaling assemblies and controlling intracellular communication(8). Although KRT17 has been shown to facilitate tumor progression in various malignancies (9), the mechanism by which KRT17 engages with oncogenic signaling pathways and whether it does so by post-transcriptionally regulating effectors that are central to the signal output remains poorly understood.

Epsin 1 (EPN1) is an endocytic adaptor protein that plays a role in membrane trafficking and signal transduction(10). Indeed, recent studies have linked members of the epsin family to cancer signaling and suggested that EPN1 may act as a functional switch between intracellular adaptor complexes and downstream transcriptional programs(11). Ubiquitin-mediated degradation is a common mechanism involved in the regulation of the stability of adaptor proteins for various signaling pathways, and E3 ligases are key factors that determine both substrate specificity and ubiquitination sites(12). However, whether KRT17 regulates the stability of EPN1 and thereby influences its role in downstream signaling has not been investigated.

In the present study, we revealed a previously unknown KRT17–EPN1 regulatory axis that modulates Wnt/β-catenin signaling in ovarian cancer cells. We show that KRT17 directly interacts with EPN1 and inhibits the SMURF1-dependent ubiquitination of EPN1 at the lysine 107 (K107) residue to stabilize EPN1. Stable EPN1 enhances Wnt/β-catenin signaling output and upregulates the expression of stemness genes. These data reveal a mechanism of crosstalk between an intermediate filament protein and Wnt/β-catenin signaling through ubiquitin-mediated regulation of an adaptor protein, which offers a fresh understanding of the regulation of cytoskeleton-associated signaling in ovarian cancer.

## 2. Material and methods

### 2.1 Patients and specimens

Between 2008 and 2013, tissues (including 143 OC specimens) were collected from patients at the Hospital of Obstetrics and Gynecology affiliated with Tongji University with the necessary informed consent. The overall survival (OS) rate was calculated from the date of surgery until the individual either passed away or reached the latest follow-up.

### 2.2 Cell Lines

The Shanghai Cell Bank of the Chinese Academy of Sciences provided all cell lines, including CAOV-3, SK-OV-3, HEK293T, IOSE80, RKO, and KGN. The cell cultures were maintained at 37°C under ambient conditions of 5% CO₂. CAOV-3 and SK-OV-3 cell lines were maintained in RPMI 1640 medium sourced from HyClone (Logan, UT, USA). HEK293T, RKO, and IOSE80 cell lines were maintained in high-glucose DMEM (enhanced with GlutaMAX™ supplement and pyruvate, sourced from Thermo Fisher Scientific). KGN cells were cultivated in DMEM/F12 medium (HyClone, Logan, Utah, United). The culture medium contained 10% fetal bovine serum (FBS), along with 100 U/ml of penicillin and 100 μg/ml of streptomycin, all obtained from Life Technologies, Carlsbad, CA, USA. Cells were dissociated and subcultured with trypsin-EDTA once they attained approximately 80% confluence, usually every three–four days. The incubation temperature was maintained at 37°C. STR analysis was performed to verify the authenticity of the cell lines.

### 2.3 Animal tumor model

Female athymic nude mice, aged six weeks, were obtained from the Slac Experimental Animal Center (Shanghai, China) and randomly distributed into various groups (n = 12). Each mouse in the groups received a subcutaneous implantation containing a total of 1 × 10⁷ SK-OV-3 OC cells, and shCtrl was compared to shKRT17 cells. Tumor volume was estimated by measuring the length and width at 8, 13, 15, 19, and 22 d after injection. Eventually, the mice were sacrificed by injecting pentobarbital sodium, and the tumors were removed for taking pictures and weighing.

### 2.4 Bioinformatics analysis

To assess the expression levels, we analyzed ovarian cancer and normal tissue samples sourced from the Gene Expression Omnibus (GEO) database. The data was sourced from GSE66957 (https://www.ncbi.nlm.nih.gov/geo/query/acc.cgi?acc = GSE66957). The dataset utilized microarray data to analyze gene expression. The methods for standardization and differential analysis were used to normalize the expression matrix using the R package affy and perform differential analysis using the R package limma. For prognosis, we selected OC samples from the Gene Expression Omnibus (GEO) database for analysis. The data was sourced from GSE140082 (https://www.ncbi.nlm.nih.gov/geo/query/acc.cgi?acc = GSE140082), and the corresponding clinical information and gene expression profiles of the cancer samples were downloaded. The dataset used gene expression microarray data, which were transformed by normalizing the expression matrix using the R package limma.

To predict the ubiquitination sites of EPN1, we accessed UniProt (https://www.uniprot.org/), searched for EPN1, and downloaded its protein sequence. We then used GPS-Uber, GPS-Uber-Ubiquitin-protein ligase enzyme, and substrate relationship prediction (biocuckoo.cn) to predict the ubiquitination modification sites of the EPN1 protein sequence.

### 2.5 Plasmid transfection and lentiviral infection

Biotechnology Limited Company in China developed lentiviral vectors targeting the desired gene to establish stable cell lines. The details of the shRNA sequences are presented in Table S1 in the Supplementary Materials. After adhering to the prescribed infection protocol and subsequent testing, shKRT17 - 2 and shKRT17 - 3 were selected to generate KRT17 knockdown CAOV-3 and SK-OV-3 cells. Negative control cell lines were created by infecting cells with lentiviruses that harbored the control shRNA. To re-express or overexpress EPN1 in OC cell lines, the target OC cells were treated with lentiviral vectors carrying complete EPN1 coding sequences or a control vector. To detect the ubiquitination sites of EPN1, HEK293T cells were transfected with site-mutant and wild-type EPN1 expression plasmids, respectively. Cells were transfected as previously described (13). To generate stable overexpression cell lines, G418 was used for screening at 350 μg/ml in CAOV-3 cells and 400 μg/ml in SK-OV-3 cells. To establish stable knockdown cell lines, puromycin was used for screening at 2 μg/mL in CAOV-3 cells and 1.5 μg/mL in SK-OV-3 cells.

### 2.6 Co - immunoprecipitation

Co-immunoprecipitation experiments were conducted as described previously (14). Initially, cell lysates containing a total of 1 mg of protein were mixed with 3 µg of specific target antibodies and incubated overnight at 4°C. Subsequently, agarose beads with protein A/G (obtained from the manufacturer) were added and incubated at 4°C for 4 h. Finally, after washing the beads, the samples were subjected to western blot analysis.

### 2.7 Immunodeficient (IHC) analysis

Tissue sections were incubated with specific primary antibodies at 4°C overnight: anti - KRT17 (A02289 - 1, BOSTER, 1:100), anti - EPN1 (14469 - 1 - AP, Proteintech, 1:100), or anti - Ki - 67 (ab16667, Abcam, 1:100). Following overnight exposure to the primary antibody, the samples were treated with a secondary antibody for 2 h at 37°C. Subsequently, the cell nuclei were stained with hematoxylin for 10 min. Subsequently, 100 μL of 3,3 - diaminobenzidine chromogenic solution (P0202, Beyotime Biotechnology, Shanghai, China) was applied to each tissue section, and the target protein expression was observed under a microscope. The staining results were scored in a single-blind manner by two independent observers. Staining intensity levels were categorized as follows: 0, no staining, 1 light intensity, 2 medium intensity; and 3, high intensity. The proportion of stained cells was evaluated using the following scale: 1 for 1–24%, 2 for 25–49%, 3 for 50–74%, and 4 for 75–100% staining. For each tumor specimen, the pair of scores was multiplied, resulting in a range of 0–12. Scores of 6 or more indicated elevated expression levels. A score of < 6 indicated minimal or absent expression.

### 2.8 Quantitative real - time PCR (qRT - PCR)

Cells underwent RNA extraction using the TRIzol reagent (Takara, Japan). Subsequently, RNA was converted into complementary DNA (cDNA) using PrimeScript™ RT Master Mix (Takara, Japan). RNA transcript levels were quantified using quantitative reverse transcription polymerase chain reaction (qRT-PCR) on the LightCycler 480 Real-Time PCR system, in conjunction with the SYBR Green PCR kit (Yeasen Biotechnology, Shanghai, China). The details of the qRT-PCR primers are provided in Table S2 in the Supplementary Document. The comparative threshold cycle (Ct) method was used to determine relative gene expression.

### 2.9 Western blot

Initially, the cell pellet was broken down with RIPA lysis buffer (Solarbio, China). Subsequently, the proteins obtained were separated using SDS-PAGE, with either a 10% or 12.5% gel. After that, the proteins were transferred to 0.45μm polyvinylidene fluoride membranes (Millipore, USA). The membranes were blocked for 2 h with 5% bovine serum albumin (BSA) at room temperature. After blocking, the membranes were exposed to primary antibodies and incubated overnight at 4°C. The following day, the membranes were incubated for 1 h with HRP-linked secondary antibodies at ambient temperature, after which they were detected using a chemiluminescent substrate (Millipore, USA). The antibodies used in this study were: anti - KRT17 (17516 - 1 - AP, Proteintech, China), anti-GAPDH (60004 - 1 - lg, Proteintech, China), anti - EPN1 (14469 - 1 - AP, Proteintech, China), anti - SOX2 (11064 - 1 - AP, Proteintech, China), anti-Nanog (14295 - 1 - AP, Proteintech, China), anti - OCT4 (60242 - 1 - Ig, Proteintech, China), anti - CD133 (A01767 - 3, Boster, China), anti - GSK3α/β (22104 - 1 - AP, Proteintech, China), anti - p - GSK3α/β (Ser21/9, 8566S, CST, USA), anti-β-catenin (51067 - 2 - AP, Proteintech, China), anti - CCND1 (ab134175, Abcam, UK), anti-β-actin (66009 - 1 - Ig, Proteintech, China), anti-ubiquitin (sc - 8017, Santa Cruz, USA), anti - SMURF1 (55175 - 1 - AP, Proteintech, China).

### 2.10 CCK8 assay

For cellular viability assays, cells were placed into 96 - well plates and then exposed to drug treatment for the specified time. Subsequently, the medium containing the drug was replaced with a new medium that included 10% CCK8 reagent. Following a 2 - h incubation period at 37°C, the absorbance was measured at 450 nm. Dimethyl sulfoxide (DMSO) was used as a negative control.

### 2.11 Colony formation assay

In 6 - well plates, each well was seeded with 2,000 cells. Subsequently, the cells were cultured under the specified conditions for 10 d. following a 10–min staining procedure with 0.2% crystal violet, the colonies were quantified.

### 2.12 Transwell migration assays

Cells were resuspended in serum-free DMEM and added to the upper section of a Transwell device, which featured a membrane with 8 μ m pores (Corning, USA) for conducting the migration assay and was provided by BD Biosciences, USA for the invasion assay. The bottom portion was supplemented with DMEM containing 10% fetal bovine serum (FBS). After 24 h, the cells that moved into the lower chamber were stained with 0.2% crystal violet.

### 2.13 Wound healing assay

An experiment was performed to evaluate the movement potential and protrusion characteristics of cancer cells using a wound healing assay. OC target cells were placed in 6 - well plates and cultured until they achieved 80 – 90% confluency. To create wounds in the cell monolayer, a sterilized one - milliliter pipette tip was employed, followed by washing away the debris using PBS. Subsequently, cell movement towards the wound area was monitored at various intervals. Cells that moved into the injury site or extended from the wound edges were observed and captured using an inverted microscope at various intervals. In each well, nine regions were randomly chosen at 100 × magnification. Cells in three wells per group were quantified in each experiment. Each experiment was performed three times, ensuring at least three repetitions.

### 2.14 Flow cytometry apoptosis assay

Cell apoptosis analysis was performed using flow cytometry with a FACS ARIA II SORP system (BD Biosciences). 1 × 10 ⁵ cells were collected and labeled with the Annexin V - FITC/PI apoptosis detection kit for 30 min, according to the manufacturer’s instructions (Beyotime, China). The samples were examined using fluorescence-activated cell sorting (FACS). Annexin V positivity combined with PI negativity marks the initial phase of apoptosis in cells. However, positivity for both Annexin V and PI indicates either advanced apoptosis or cell necrosis.

### 2.15 Immunoprecipitation - Mass Spectrometry (IP - MS)

Co-IP assays were conducted as previously described, followed by the steps mentioned below. For protein digestion, DTT solution was added to a final concentration of 10 mmol/L and incubated in a 56 ° C water bath for 1 h. Follow this by adding **iodoacetamide** to a concentration of 55 mmol/L, and the reaction was allowed to proceed in the dark for 40 min. Trypsin was administered and allowed to incubate at 37° C overnight. After digestion, the peptides were desalted with a self-priming column and concentrate by 45°C vacuum centrifugation at 45°C. A 1% formic acid solution was prepared, mixed using a vortex, centrifuged at 13200 rpm for 10 min at 4 °C, and the supernatant was used for mass spectrometry analysis. For Nano LC-MS/MS analysis, the ThermoFisher Easy-nLC 1200 system with designated nano-columns was used, and a 5 μL volume was injected. Mobile phase A was a 0.1% formic acid aqueous solution, and B was a mixture of 0.1% formic acid and 80% acetonitrile, with a flow rate of 600 nL/min and a specific gradient. The Q Exactive™ mass spectrometer was employed with a 2.2 kV spray voltage and 270 ° C capillary temperature. Specific MS1 and MS2 parameters were set, and the top 20 most intense peptide ions were selected for data-dependent MS2 analysis. MaxQuant (1.6.2.10) was used to analyze raw MS files against the target database according to the sample species. The parameters were set as follows: fixed carbamidomethylation (C), variable oxidation (M) and acetylation (N - term), trypsin specificity, up to two missed cleavage sites, and a 20 ppm mass tolerance for MS1 and MS2. For downstream analysis, only high-confidence peptides were used.

### 2.16 Protein degradation and cellular ubiquitination assay

To assess EPN1 protein degradation, cells were treated with cycloheximide (CHX, 100 μg/ml, MCE, USA) at different time intervals before collection for immunoblot analysis. To assess the ubiquitination levels of EPN1, cells were transfected with Flag-Ub plasmid and collected 48 h post-transfection. Prior to collection, the cells were exposed to 10 μ M MG132 for 4 h. Protein extracts were obtained using SDS lysis buffer composed of 50 mM Tris-HCl at pH 7.5. The samples were treated with a solution containing 1% SDS and 10 mM DTT, followed by heating at 95°C for ten minutes. The lysates were diluted tenfold using gentle lysis buffer before immunoprecipitation and immunoblotting.

### 2.17 Spheroid formation assay

OC cell single-cell suspensions with a concentration of 4 × 10² cells/mL were placed into 24 - well ultra - low attachment plates (Corning, USA). Subsequently, these cells were cultured in DMEM/F12 medium without phenol red (Gibco). The medium was enriched with B27 (Gibco, USA, #12587010), 20 ng/mL epidermal growth factor (EGF; Solarbio, China, #P00033), 20 ng/mL basic fibroblast growth factor (bFGF; Solarbio, China, #P00032), and 5 μg/mL insulin (Solarbio, China, #I8040). Tumorspheres were examined using a phase-contrast microscope (Olympus, IX53 + DP80).

### 2.18 Aldehyde dehydrogenase (ALDH) activity assays

ALDH activity was detected using the ALDH Activity Assay Kit (Boxbio Science & Technology) according to the manufacturer’s protocol. Briefly, the cells were collected by centrifugation in centrifuge tubes. Samples were prepared by maintaining a ratio between the number of cells (10,000 cells) and the amount of extraction solution (in mL), ranging from 500 to 1000. The specimens were subsequently subjected to sonication on ice (300 W power, 3 s of sonication, with 7 - s intervals, for a total duration of 3 min). Subsequently, the samples were centrifuged at 15,000 × g for 20 min at 4°C. The supernatant was transferred to a 96 - well UV plate, followed by the addition of the assay reagent. A microplate reader was used to measure the absorbance of the samples at 340 nm to determine ALDH activity.

### 2.19 Statistical analysis

For each experiment, a minimum of three trials were conducted. Unless indicated otherwise, the data are presented as the mean ± standard deviation (SD). SPSS Statistics (version 20.0) and GraphPad (version 8.0.2) were utilized to analyze the data. Statistical comparisons between the control and experimental groups were conducted using either two-tailed Student’s t-tests or one-way ANOVA. A two-way ANOVA was employed to compare the growth curves between the two groups. The Mann - Whitney U test was applied to examine the relationship between the expression of KRT17 or EPN1 and clinicopathological features. Spearman’s correlation was used to assess the association between variables. The Kaplan–Meier method was used for survival analysis. A significance level of 0.05 was used to assess the statistical relevance.

The original images of the western blots can be found in the Original WB blots section of the Supplementary Materials.

## 3. Results

### 3.1 KRT17 is upregulated in ovarian cancer and correlates with poor patient survival

To identify genes potentially involved in OC progression, we compared transcriptomic profiles between OC tissues and normal ovarian tissues using GEO datasets. To identify robustly dysregulated and clinically relevant genes, we set stringent filtering thresholds: log₂FoldChange > 1.5 and adjusted P < 0.001 for differential expression, and hazard ratio (HR) > 1.5 (P < 0.05) for survival correlation. Five genes—FAM117B, KAZALD1, KRT17, LPHN2, and ZNF608—were selected, as their higher expression levels were associated with poorer OS in OC patients. Among these candidates, KRT17 displayed the highest basal expression in SK-OV-3 cells (Fig. 1a).

**Fig. 1.**
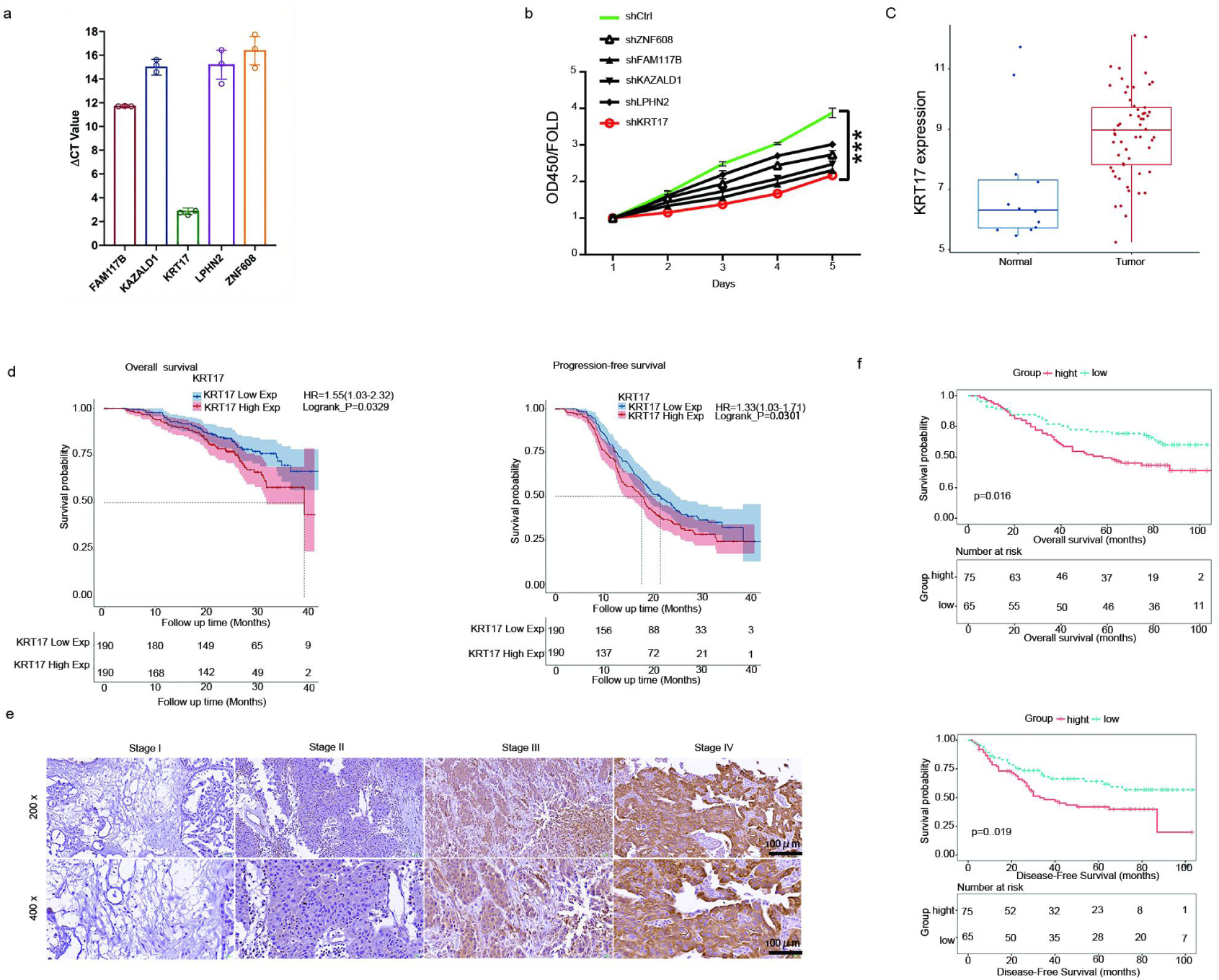
Higher KRT17 levels track with poorer outcomes in ovarian cancer. a. Baseline mRNA levels of FAM117B, KAZALD1, KRT17, LPHN2, and ZNF608 in SK-OV-3 cells (qRT–PCR; higher ΔCt indicates lower expression). b. SK-OV-3 cell proliferation after silencing individual candidate genes (CCK-8). c. KRT17 mRNA levels in ovarian cancer and normal ovarian tissues from GEO datasets. d. Kaplan–Meier plots of overall survival (OS) and progression-free survival (PFS) grouped by KRT17 expression in GEO cohorts. e. IHC staining of KRT17 across ovarian cancer stages (I–IV). f. OS and disease-free survival (DFS) curves according to KRT17 protein levels in ovarian cancer patients. ***P < 0.001.

To functionally prioritize these genes, SK-OV-3 cells were infected with lentiviral shRNAs targeting each candidate. CCK-8 assays showed that KRT17 depletion produced the strongest inhibitory effect on cell proliferation compared with knockdown of the other genes (Fig. 1b). Consistent with these functional screening results, GEO dataset analysis confirmed that KRT17 expression was significantly higher in OC tissues than in normal ovarian tissues (adjusted P = 0.000868), with a fold change of 1.81 (Fig. 1c).

For prognostic evaluation, patients were separated into high- and low-expression groups using the median KRT17 level as a cutoff. Kaplan–Meier curves revealed that patients with elevated KRT17 expression experienced shorter OS (log-rank P = 0.0329, HR = 1.55, 95% confidence interval [CI] 1.03–2.32) and progression-free survival (PFS) (log-rank P = 0.0301, HR = 1.33, 95% CI 1.03–1.71) (Fig. 1d). This expression pattern was further supported at the protein level, as immunohistochemical staining demonstrated a stage-associated increase in KRT17 expression across ovarian cancer stages I–IV (Fig. 1e, Table 1 and 2). Consistently, clinicopathological correlation analyses indicated that KRT17 expression was associated with tumor size and metastatic status (Table 1 and 2). In addition, tissue microarray-based survival analysis confirmed that higher KRT17 protein abundance correlated with reduced OS and disease-free survival (DFS) in OC patients (Fig. 1f).

**Table 1.** Relationship between KRT17 expression and tumor characteristics in patients with ovarian cancer.

| Features | No. of patients | KRT17 expression |  | p value |
| --- | --- | --- | --- | --- |
|  |  | low | high |  |
| All patients | 140 | 65 | 75 |  |
| Age* |  |  |  | 0.990 |
| ≤51years | 71 | 33 | 38 |  |
| > 51years | 69 | 32 | 37 |  |
| Stage |  |  |  | <u>0.017</u> |
| I | 12 | 8 | 4 |  |
| II | 34 | 18 | 16 |  |
| III | 68 | 32 | 36 |  |
| IV | 26 | 7 | 19 |  |
| Tumor size** |  |  |  | <u>0.011</u> |
| ≤11.95 cm | 70 | 40 | 30 |  |
| > 11.95 cm | 70 | 25 | 45 |  |
| T Infiltrate |  |  |  | 0.074 |
| T1 | 12 | 8 | 4 |  |
| T2 | 34 | 18 | 16 |  |
| T3 | 94 | 39 | 55 |  |
| Lymphatic metastasis (N) |  |  |  | 0.356 |
| N0 | 107 | 52 | 55 |  |
| N1 | 33 | 13 | 20 |  |
| Metastasis |  |  |  | <u>0.028</u> |
| M0 | 114 | 58 | 56 |  |
| M1 | 26 | 7 | 19 |  |
| Relapsing state |  |  |  | 0.206 |
| No | 30 | 17 | 13 |  |
| Yes | 110 | 48 | 62 |  |

TNM Eighth Edition of tumor-node-metastasis classification. The underlined values were considered statistically significant. \*51 years old is the median age of all ovarian cancer patients; \*\*11.95 cm is the median tumor size of all ovarian cancer patients.

**Table 2.** Relationship between KRT17 expression and tumor characteristics in patients with ovarian cancer.

|  |  | **KRT17** |
| --- | --- | --- |
| **Tumor size** | Spearman correlation | 0.215 |
|  | Sig. (two-tailed) | <u>0.011</u> |
|  | N | 140 |
| **Metastasis** | Spearman correlation | 0.202 |
|  | Sig. (two-tailed) | <u>0.016</u> |
|  | N | 140 |
| **Stage** | Spearman correlation | 0.187 |
|  | Sig. (two-tailed) | <u>0.027</u> |
|  | N | 140 |

TNM Eighth Edition of tumor-node-metastasis classification. The underlined values were considered statistically significant.

### 3.2 KRT17 knockdown suppresses malignant phenotypes of ovarian cancer cells in vitro and in vivo

We next examined endogenous KRT17 expression in normal ovarian epithelial cells (IOSE80) and several OC cell lines. qRT-PCR analysis showed that KRT17 was highly expressed in SK-OV-3 and CAOV-3 cells, whereas KGN cells exhibited relatively lower expression compared with IOSE80 cells (Fig. 2a). Based on these results, SK-OV-3 and CAOV-3 cells were selected for subsequent loss-of-function experiments.

**Fig. 2.**
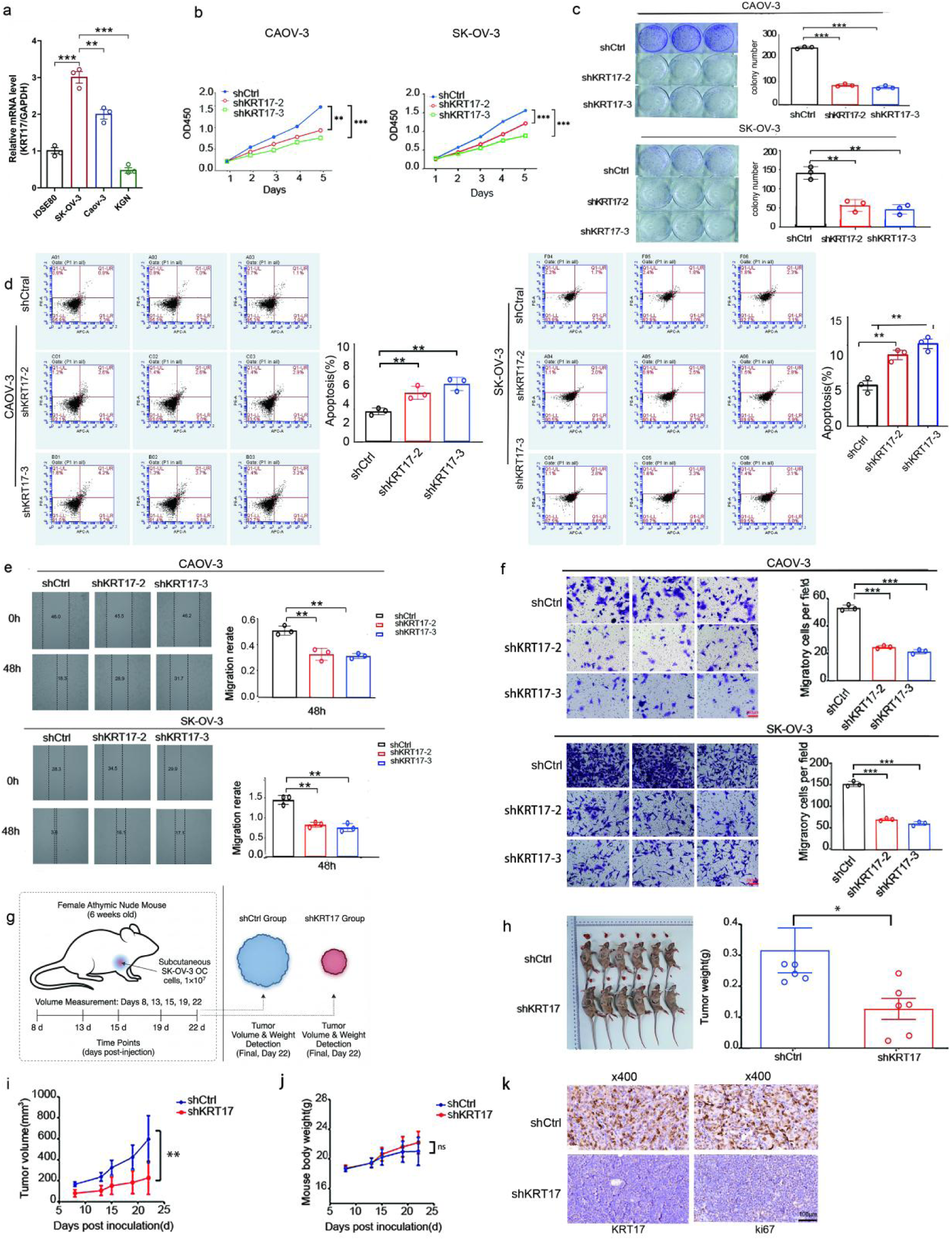
Silencing KRT17 limits ovarian cancer cell growth and migration and increases apoptosis. a. KRT17 mRNA levels across normal ovarian epithelial cells and ovarian cancer cell lines (qRT–PCR). b–d. Growth and apoptosis changes following KRT17 knockdown in CAOV-3 and SK-OV-3 cells measured by CCK-8 (b), colony formation ©, and flow cytometry (d). e–f. Migration after KRT17 knockdown examined by wound-healing (e) and Transwell assays (f). g. Outline of the xenograft experiment. h–i. Xenograft tumors formed by SK-OV-3 cells at day 22: gross appearance (h) and tumor weight (i). j. Mouse body weight recorded during tumor growth. k. IHC staining of KRT17 and Ki-67 in xenograft tumors. *P < 0.05; **P < 0.01; ***P < 0.001.

Three independent shRNAs targeting KRT17 were generated, and shKRT17-2 and shKRT17-3, which achieved the most efficient knockdown, were chosen for subsequent assays (Supplementary Fig. 1). Stable KRT17-silenced SK-OV-3 and CAOV-3 cell lines were established, and the knockdown efficiency was validated at both the mRNA and protein levels (Supplementary Fig. 2).

Functional characterization showed that KRT17 silencing reduced cell growth and clonogenic potential, as assessed by CCK-8 and colony formation assays (Fig. 2b–c). Flow cytometry further indicated that the proportion of apoptotic cells increased after KRT17 knockdown in both cell lines (Fig. 2d). Moreover, wound-healing and transwell assays consistently demonstrated that cell migration was impaired upon KRT17 depletion (Fig. 2e–f).

To evaluate in vivo relevance, SK-OV-3 cells stably expressing shCtrl or shKRT17 were implanted into nude mice. Tumors derived from shKRT17-expressing cells showed markedly reduced tumor volume and weight relative to controls (Fig. 2g–i), whereas mouse body weight remained comparable between groups (Fig. 2j). Immunohistochemical staining of the xenograft tumors further revealed lower KRT17 and Ki-67 expression in the shKRT17 group (Fig. 2k).

### 3.3 KRT17 binds EPN1 and prevents SMURF1-mediated ubiquitination at K107, thereby stabilizing EPN1

To investigate the mechanism by which KRT17 exerts its pro-tumorigenic effects, we performed immunoprecipitation followed by mass spectrometry (IP–MS) to identify proteins associated with KRT17 in ovarian cancer cells. By integrating IP–MS candidates with UbiBrowser 2.0 and DegPred predictions, EPN1 emerged as the top-ranked interacting protein, and SMURF1 was predicted as a potential E3 ligase targeting EPN1 (interaction score = 37.019).

Co-immunoprecipitation assays validated the interaction between KRT17 and EPN1 (Fig. 3a) and confirmed the binding between SMURF1 and EPN1 (Fig. 3b). Notably,a reduction in KRT17 resulted in a marked increase in the SMURF1-EPN1 interaction (Fig. 3c), indicating that KRT17 negatively regulates their association. When KRT17 was depleted, EPN1 protein levels decreased substantially, whereas EPN1 mRNA levels remained largely unchanged in both CAOV-3 and SK-OV-3 cells (Fig. 3d–e), indicating regulation at the post-transcriptional level.

**Fig. 3.**
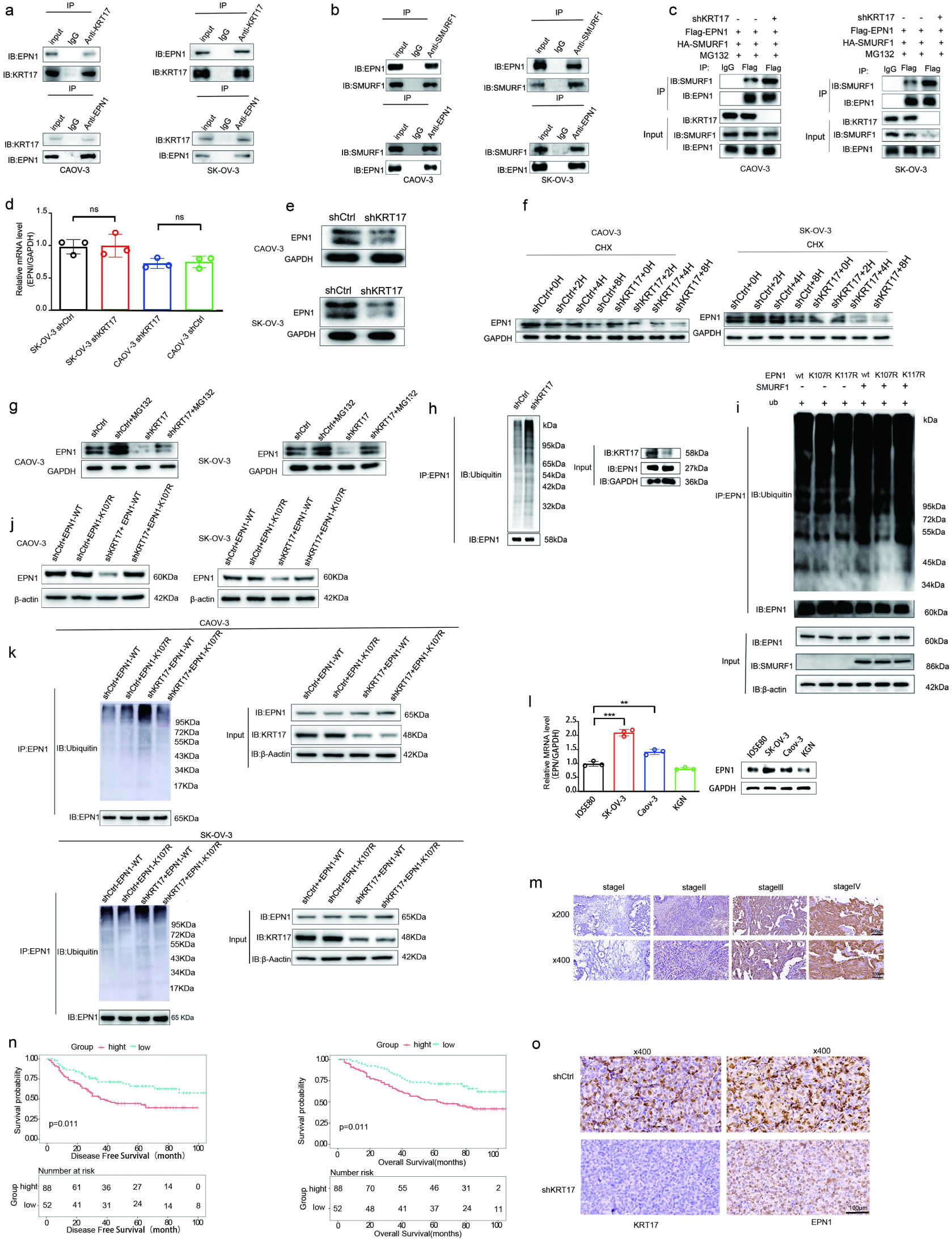
KRT17 maintains EPN1 stability by limiting SMURF1-dependent ubiquitination. a. Co-IP confirming interaction between KRT17 and EPN1 in CAOV-3 and SK-OV-3 cells. b. Co-IP showing binding between SMURF1 and EPN1. c. SMURF1–EPN1 interaction in cells expressing shCtrl or shKRT17. d–e. EPN1 transcript (d) and protein (e) levels after KRT17 knockdown (qRT–PCR and Western blot). f. Cycloheximide chase analysis of EPN1 protein turnover. g. EPN1 protein levels after MG132 treatment. h. EPN1 ubiquitination in shCtrl and shKRT17 cells detected by immunoprecipitation followed by ubiquitin immunoblotting. i. SMURF1-driven ubiquitination of EPN1-WT and EPN1 mutants (K107R, K117R). j–k. EPN1 protein abundance (j) and ubiquitination status (k) under the indicated conditions. l. EPN1 expression in ovarian cancer cell lines (qRT–PCR and Western blot). m. EPN1 staining in ovarian cancer tissues at different stages (IHC). n. Survival curves (OS and PFS) plotted by EPN1 expression level. o. IHC staining of EPN1 and Ki-67 in xenograft tumors.

Cycloheximide (CHX) chase assays, which block de novo protein synthesis, further showed that EPN1 turnover was markedly accelerated in KRT17-silenced cells (Fig. 3f). In line with proteasome-dependent degradation, treatment of proteasome inhibitor MG132 restored EPN1 protein abundance following KRT17 knockdown (Fig. 3g). Consistently, ubiquitination assays demonstrated that polyubiquitinated EPN1 accumulated to higher levels upon KRT17 depletion (Fig. 3h).

To map the SMURF1-dependent ubiquitination site(s) on EPN1, lysine residues K107 and K117 were individually mutated to arginine. Ubiquitination assays showed that the K107R mutation markedly reduced SMURF1-mediated ubiquitination, whereas the K117R mutation had little effect on it (Fig. 3i). Reconstitution experiments further supported the importance of K107: in KRT17-depleted CAOV-3 and SK-OV-3 cells, wild-type EPN1 displayed reduced stability and increased ubiquitination, while the K107R mutant remained stable and was largely resistant to ubiquitination changes induced by KRT17 knockdown (Fig. 3j–k).

The clinical relevance of EPN1 was also evaluated. EPN1 protein levels were markedly elevated in epithelial ovarian cancer cell lines (SK-OV-3 and Caov-3) compared with the non-tumorigenic ovarian epithelial cell line IOSE80, while lower EPN1 expression was also observed in the human ovarian granulosa cell tumor cell line KGN (Fig. 3l). Immunohistochemical staining of a tissue microarray revealed a stage-associated increase in EPN1 protein expression across ovarian cancer tissues (Fig. 3m, Table 3 and 4). Consistently, higher EPN1 expression was associated with more advanced clinicopathological characteristics, including lymphatic metastasis and metastatic status (Table 3 and 4). Kaplan–Meier analysis further indicated that patients with high EPN1 expression had shorter OS and DFS compared with those with low expression (Fig. 3n). In addition, xenograft tumors derived from KRT17-knockdown cells exhibited reduced EPN1 protein levels, as confirmed by immunohistochemical staining (Fig. 3o).

**Table 3.** Relationship between EPN1 expression and tumor characteristics in patients with ovarian cancer.

| Features | No. of patients | EPN1 expression |  | p value |
| --- | --- | --- | --- | --- |
|  |  | low | high |  |
| All patients | 140 | 52 | 88 |  |
| Age* |  |  |  | 0.050 |
| ≤51years | 71 | 32 | 39 |  |
| > 51years | 69 | 20 | 49 |  |
| Tumor size** |  |  |  | 0.486 |
| ≤11.95 cm | 70 | 28 | 42 |  |
| > 11.95 cm | 70 | 24 | 46 |  |
| T |  |  |  | 0.052 |
| T1 | 12 | 7 | 5 |  |
| T2 | 34 | 15 | 19 |  |
| T3 | 94 | 30 | 64 |  |
| Lymphatic metastasis (N) |  |  |  | <u>0.003</u> |
| N0 | 107 | 47 | 60 |  |
| N1 | 33 | 5 | 28 |  |
| Metastasis |  |  |  | <u>0.003</u> |
| M0 | 114 | 49 | 65 |  |
| M1 | 26 | 3 | 23 |  |
| Stage |  |  |  | <u>0.004</u> |
| I | 12 | 7 | 5 |  |
| II | 34 | 15 | 19 |  |
| III | 68 | 27 | 41 |  |
| IV | 26 | 3 | 23 |  |

TNM Eighth Edition of tumor-node-metastasis classification. The underlined values were considered statistically significant. \*51 years old is the median age of all ovarian cancer patients; \*\*11.95 cm is the median tumor size of all ovarian cancer patients.

**Table 4.** Relationship between KRT17 expression and tumor characteristics in patients with ovarian cancer.

|  |  | EPN1 |
| --- | --- | --- |
| **Lymphatic metastasis (N)** | Spearman correlation | 0.253 |
|  | Sig. (two-tailed) | <u>0.003</u> |
|  | N | 140 |
| **Metastasis** | Spearman correlation | 0.253 |
|  | Sig. (two-tailed) | <u>0.003</u> |
|  | N | 140 |
| **Stage** | Spearman correlation | 0.245 |
|  | Sig. (two-tailed) | <u>0.004</u> |
|  | N | 140 |

TNM Eighth Edition of tumor-node-metastasis classification. The underlined values were considered statistically significant.

### 3.4 EPN1 mediates the effects of KRT17 on ovarian cancer cell proliferation and **migration**

To clarify the functional relationship between KRT17 and EPN1, rescue experiments were carried out in CAOV-3 and SK-OV-3 cells. Cells were transfected with an EPN1 overexpression construct alone or together with shKRT17, and successful expression was confirmed at both the mRNA and protein levels (Supplementary Fig. 3).

EPN1 re-expression counteracted the growth-inhibitory and migration-suppressive effects of KRT17 knockdown. Specifically, CCK-8, colony formation, and wound-healing assays showed that EPN1 overexpression restored cell proliferation, clonogenicity, and migratory capacity in KRT17-depleted cells (Fig. 4a–c).

**Fig. 4.**
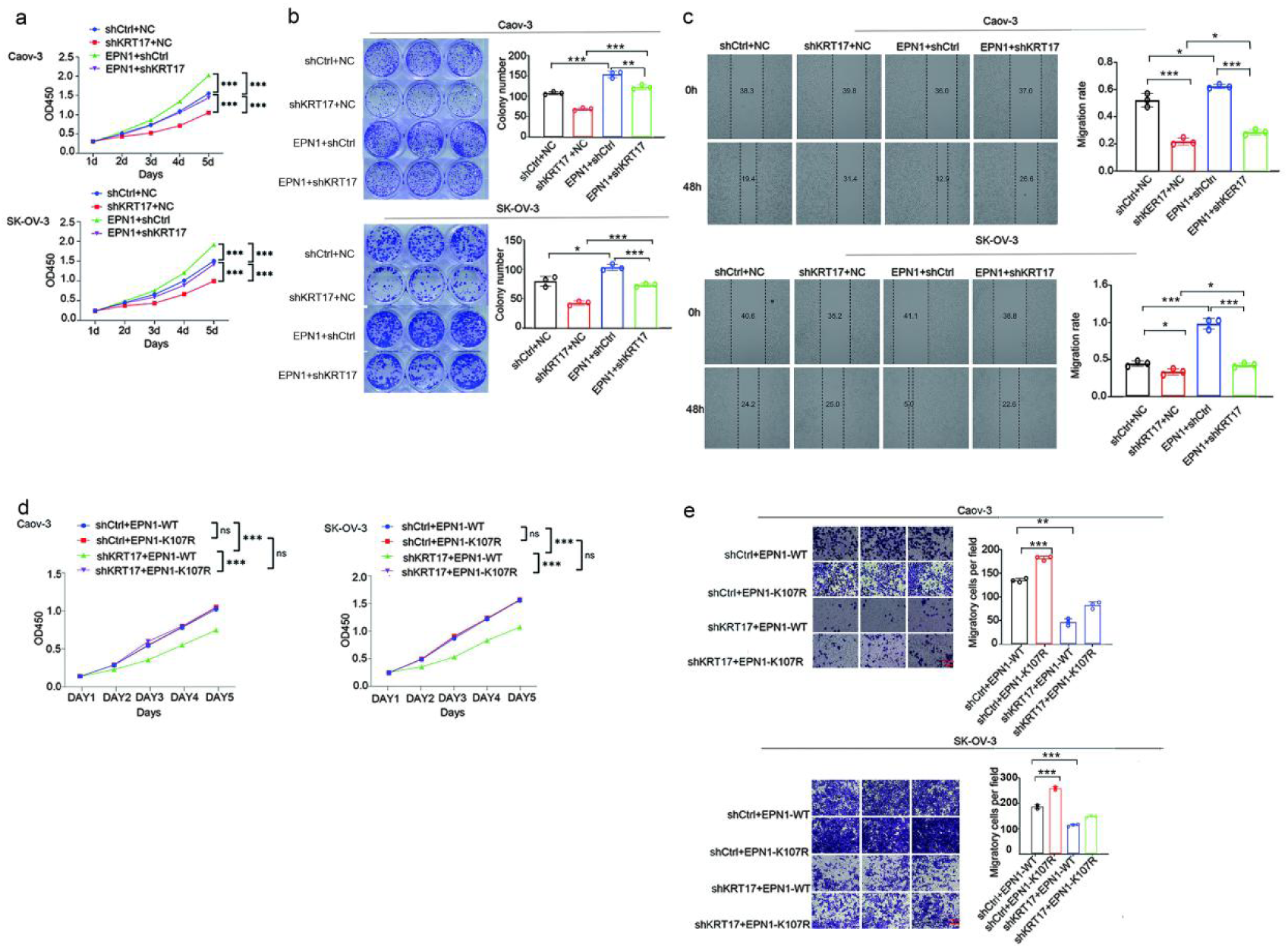
EPN1 mediates the effects of KRT17 on proliferation and migration. a–c. Proliferation and migration in the indicated groups measured by CCK-8 (a), colony formation (b), and wound-healing assays ©. d–e. CCK-8 (d) and Transwell migration (e) results for cells expressing EPN1-WT or EPN1-K107R, with or without KRT17 knockdown. *P < 0.05; **P < 0.01; ***P < 0.001; ns, not significant.

To determine whether this rescue depended on EPN1 stability, either wild-type EPN1 or ubiquitination-resistant mutant EPN1-K107R was introduced into KRT17-silenced cells. In cells expressing wild-type EPN1, the suppressive effects of KRT17 knockdown on proliferation and migration remained evident. In contrast, EPN1-K107R largely recovered proliferative and migratory abilities, supporting the idea that KRT17 influences malignant phenotypes through ubiquitination-dependent regulation of EPN1 stability (Fig. 4d–e).

### 3.5 The KRT17–EPN1 axis modulates Wnt/β-catenin signaling and cancer stemness

Given the reported involvement of EPN1 in Wnt/β-catenin signaling(15), we next examined whether the KRT17–EPN1 axis affects this pathway. Western blot analysis showed that KRT17 knockdown resulted in reduced levels of phosphorylated GSK3α/β, β-catenin, and CCND1, together with decreased expression of stemness-associated markers, including SOX2, OCT4, CD133, and NANOG, in CAOV-3 and SK-OV-3 cells (Fig. 5a–b). Similar trends were observed in xenograft tumors derived from KRT17-silenced cells (Fig. 5c–d).

**Fig. 5.**
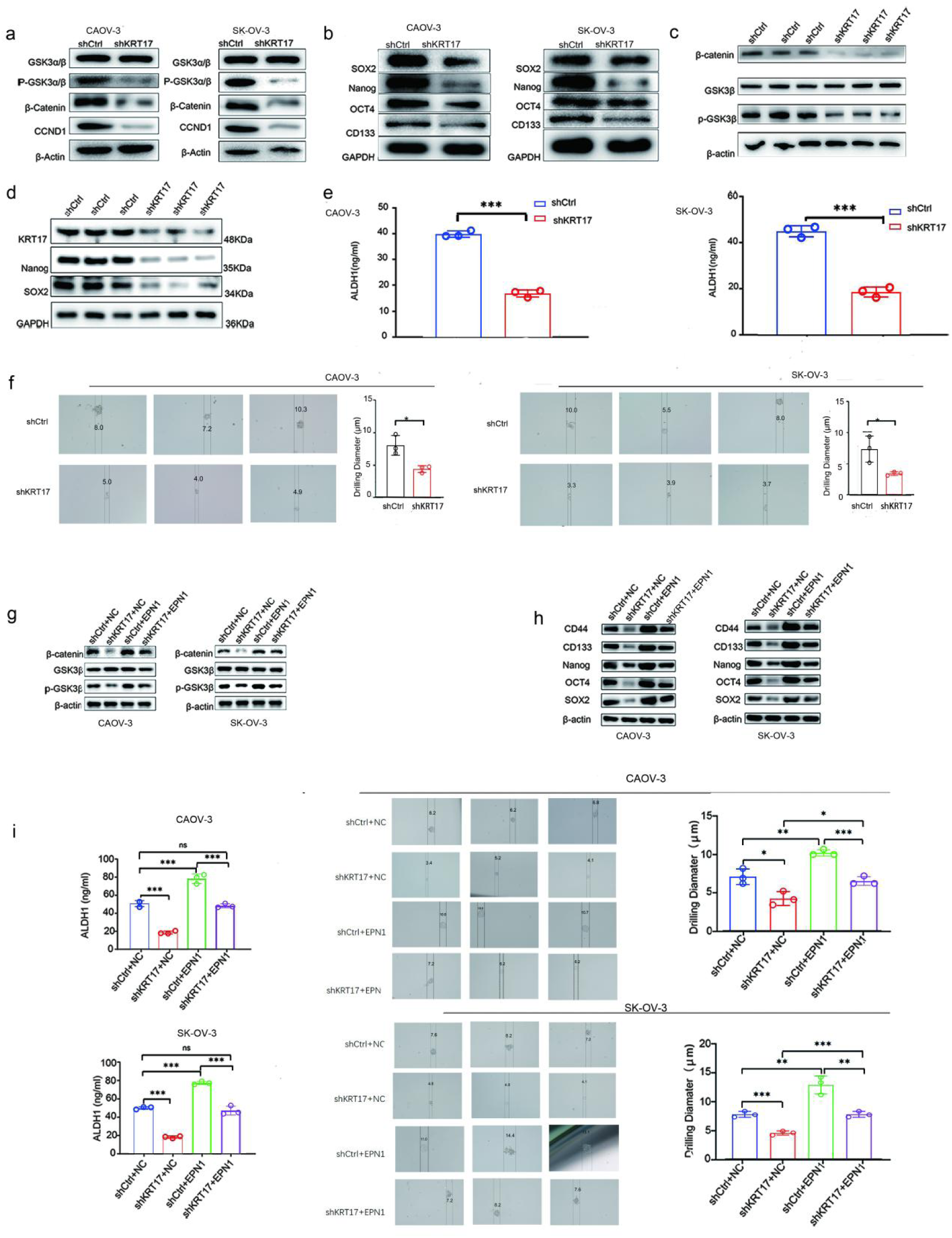
KRT17–EPN1 signaling shapes Wnt/β-catenin activity and stem-like traits. a–b. Wnt/β-catenin pathway proteins (a) and stemness markers (b) in shCtrl and shKRT17 cells (Western blot). c–d. Wnt/β-catenin signaling proteins (c) and stemness markers (d) in xenograft tumors. e–f. Stem-like features measured by ALDH activity (e) and tumorsphere formation (f). g–j. EPN1 re-expression restores Wnt/β-catenin signaling (g), stemness markers (h), ALDH activity (i), and tumorsphere formation (j).

Functional assays further supported the reduction in stem-like properties following KRT17 depletion. Specifically, ALDH1 activity decreased and tumorsphere formation was impaired in KRT17-knockdown cells (Fig. 5e–f). Importantly, re-expression of EPN1 in these cells restored the levels of phosphorylated GSK3α/β, β-catenin abundance, stemness marker expression, ALDH1 activity, and tumorsphere-forming capacity (Fig. 5g–j).

### 3.6. The KRT17–EPN1 axis influences cisplatin response and chemoresistance

We also assessed whether this signaling axis influenced the chemotherapeutic response. KRT17 depletion reduced basal proliferation and enhanced cisplatin-mediated growth inhibition. In both CAOV-3 and SK-OV-3 cells, silencing KRT17 led to a clear increase in cisplatin sensitivity. This was reflected by a pronounced reduction in cell viability in CCK-8 assays, a stronger growth-inhibitory effect of cisplatin, and markedly decreased clonogenic survival after drug treatment (Fig. 6a–b).

**Fig. 6.**
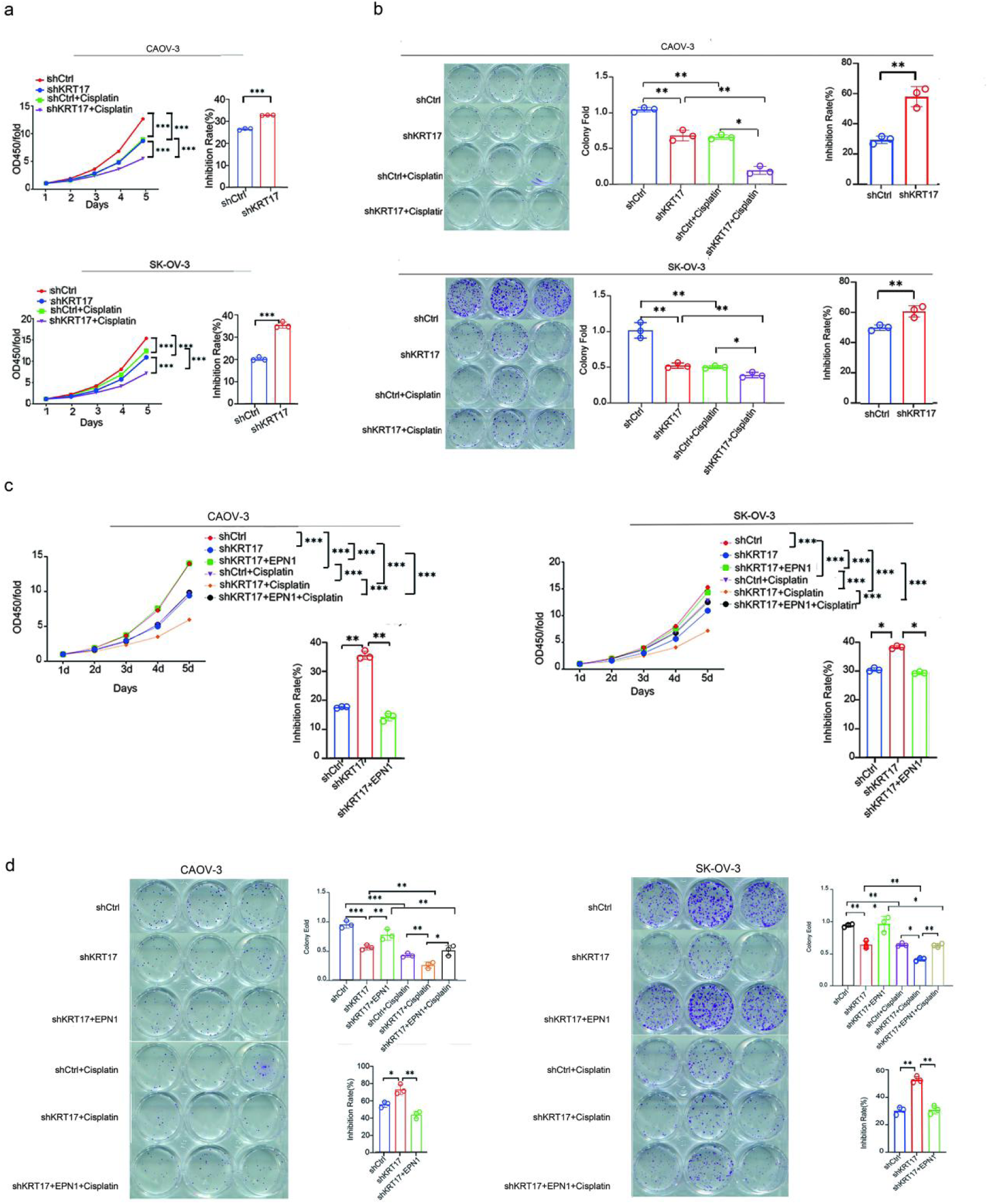
KRT17–EPN1 signaling alters the cisplatin response in ovarian cancer cells. a–b. Cisplatin (10 μM) response measured by CCK-8 (a) and colony formation (b) assays. c–d. Impact of EPN1 re-expression on cisplatin (10 μM) response assessed by CCK-8 (c) and colony formation (d).

When EPN1 expression was restored in KRT17-depleted cells, the enhanced inhibitory effect of cisplatin was partially reversed. Re-expression of EPN1 reduced cisplatin-induced growth inhibition and restored, at least in part, cell viability and colony-forming capacity in the presence of cisplatin (Fig. 6c–d). These observations suggest that EPN1 plays a key role in mediating the altered cisplatin response caused by KRT17 loss.

Collectively, these results point to a contribution of the KRT17–EPN1 axis to cisplatin resistance in ovarian cancer cells.

## Discussion

In this study, we describe a cytoskeleton-associated regulatory mechanism in which KRT17 modulates Wnt/β-catenin–associated signaling outputs by stabilizing the adaptor protein EPN1 (Fig. 7). Our data support a model in which KRT17 physically associates with EPN1 and weakens SMURF1-driven ubiquitination at lysine 107 (K107), thereby limiting the proteasomal turnover of EPN1. By preserving EPN1 abundance, KRT17 is associated with sustained Wnt/β-catenin pathway engagement, linking an intermediate filament protein to a core oncogenic signaling axis in ovarian cancer and providing mechanistic insight into how intracellular communication may be shaped in this disease. Intermediate filament proteins have traditionally been viewed as structural elements that safeguard cellular architecture (30). However, increasing evidence indicates that cytoskeletal components can participate more directly in signaling by acting as scaffolds or regulatory platforms (31–33). Our findings support this emerging view: rather than serving only as a passive structural marker, KRT17 appears to influence signal transmission. The KRT17–EPN1 interaction offers a tangible route through which cytoskeletal organization can shape downstream signaling behavior, underscoring a mode of cytoskeleton–signaling crosstalk that may contribute to oncogenic pathway engagement.

**Fig. 7.**
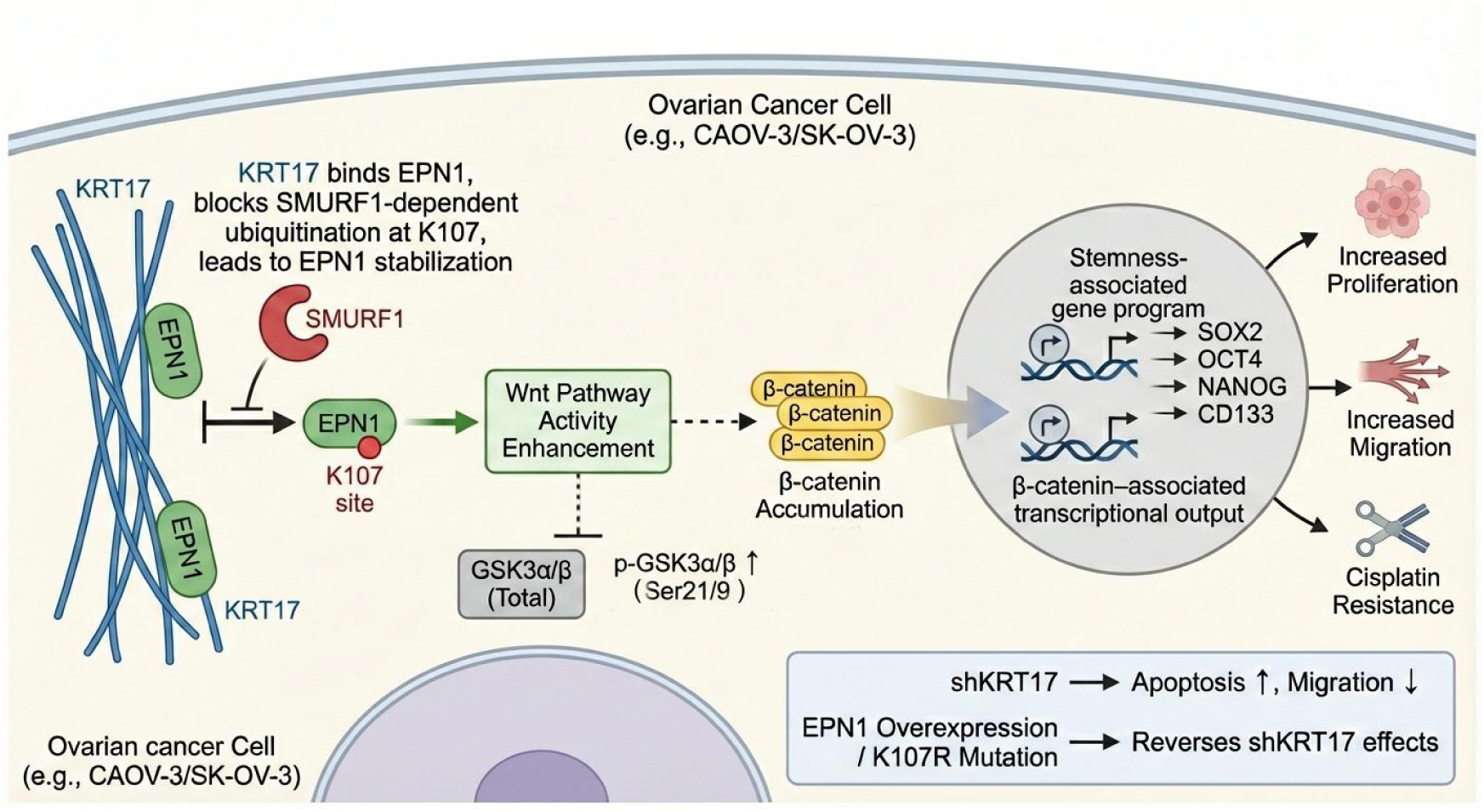
Model of the KRT17–EPN1 signaling axis. KRT17 binds EPN1 and reduces SMURF1-dependent ubiquitination at K107, helping maintain EPN1 stability. Stabilized EPN1 supports Wnt/β-catenin activity, reinforcing cell growth, migration, stem-like traits, and cisplatin resistance in ovarian cancer cells.

Our results also broaden the functional picture of EPN1 in cancer-related signaling. Beyond its well-established role in endocytosis, EPN1 behaves as a regulated signaling node whose abundance affects downstream pathway outputs. We identified SMURF1 as an E3 ubiquitin ligase that promotes EPN1 ubiquitination and showed that modification at K107 is a key determinant of EPN1 stability. In this setting, binding of KRT17 reduces ubiquitination and thereby maintains EPN1 protein levels. More generally, these data emphasize how post-translational regulation, particularly ubiquitination, can modulate adaptor protein abundance and, consequently, signaling strength.

A major downstream consequence of KRT17-dependent EPN1 stabilization is the alteration of Wnt/β-catenin associated signaling readouts. In our study, KRT17 depletion reduced phosphorylated GSK3α/β, total β-catenin, and the downstream effector CCND1, accompanied by decreased expression of stemness-associated markers in ovarian cancer cells and xenograft tumors. Notably, we primarily assessed total β-catenin abundance rather than its subcellular localization; therefore, nuclear β-catenin accumulation and direct transcriptional activity were not evaluated. Nevertheless, coordinated changes in total β-catenin and CCND1, together with the EPN1 rescue experiments, support a functional link between the KRT17–EPN1 module and Wnt/β-catenin pathway output in this context.

Sustained Wnt/β-catenin signaling has been extensively implicated in the maintenance of stem-like tumor phenotypes (34–35). Although we did not directly test the requirement of Wnt/β-catenin signaling for stemness using pathway-specific inhibitors or genetic blockade, modulation of the KRT17–EPN1 axis consistently altered stemness-associated marker expression, ALDH1 enzymatic activity, and tumorsphere-forming capacity. These observations are therefore consistent with established models in which β-catenin–associated signaling contributes to cancer cell plasticity and self-renewal.

In parallel, engagement of the KRT17–EPN1 axis was associated with reduced sensitivity to cisplatin, whereas KRT17 depletion enhanced chemotherapeutic responsiveness. Importantly, re-expression of EPN1 partially reversed cisplatin sensitivity, restoring cell viability and clonogenic potential in the presence of drug treatment. These findings suggest that altered cisplatin response is likely an indirect consequence of signaling-dependent shifts toward a stem-like cellular state rather than a direct effect on drug transport or metabolism, aligning with the concept that chemoresistance can emerge from pathway-driven cellular programs (36–37).

Several limitations should be acknowledged. Although SMURF1 was identified as a key E3 ligase regulating EPN1 ubiquitination in our models, other ubiquitin-related enzymes may also participate in EPN1 turnover in a context-dependent manner. Future systematic analyses are required to define the broader ubiquitin network involved in this process. In addition, our mechanistic findings were primarily based on ovarian cancer cell lines and xenograft models, which may not fully reflect tumor heterogeneity. Thus, the strength and prevalence of the KRT17–EPN1 axis may vary across tumor subtypes or cancer types, a point that warrants further investigation but does not alter the core mechanistic framework proposed here.

In summary, we propose that KRT17 preserves EPN1 by limiting SMURF1-dependent ubiquitination at K107, and that this stabilization is associated with sustained Wnt/β-catenin signaling output, enhanced stem-like properties, and reduced chemotherapeutic sensitivity. These findings expand the view of intermediate filaments from structural elements to active contributors to signaling control and illustrate how post-translational regulation of adaptor proteins can shape oncogenic programs and cellular plasticity in ovarian cancer.

## List of abbreviations

OC: ovarian cancer
ALDH: aldehyde dehydrogenase
BSA: bovine serum albumin
CCK-8: cell counting kit-8
CHX: cycloheximide
EMT: epithelial - mesenchymal transition
DFS: disease-free survival
DMEM: Dulbecco’s modified Eagle medium
DMSO: dimethyl sulfoxide
DTT: dithiothreitol
E3: E3 ubiquitin ligase
EPN1: epsin 1
FACS: fluorescence-activated cell sorting
FBS: fetal bovine serum
GEO: gene expression omnibus
HR: hazard ratio
IHC: immunohistochemistry
IP–MS: immunoprecipitation–mass spectrometry
K107: lysine 107
KRT17: keratin 17 OS - overall survival
PFS: progression-free survival
PI: propidium iodide
qRT-PCR: quantitative real-time polymerase chain reaction
SD: standard deviation
SMURF1: SMAD ubiquitination regulatory factor 1
CHX: Cycloheximide
Ub: ubiquitin

## Declarations

### Ethics approval and consent to participate

All animal experiments were approved by the Animal Care and Use Committee of the Tongji University. All animal experiments were conducted in accordance with the ARRIVE guidelines 2.0.The animal experiment ethics certificate number is TJBG10321201. All clinical samples were obtained with informed consent from the patients before collection. The Ethics Committee at the Tongji University - affiliated Hospital of Obstetrics and Gynecology granted approval for the analysis of the patient samples.The medical research ethics certificate number is KS25118.

### Consent for publication

All authors were consent for publication.

## Acknowledgements

Not applicable.

## Data availability

Data availability: The original data used and analyzed in this study are available from the corresponding author upon reasonable requests.

## CRediT authorship contribution statement

Guiqiang Du: Conceptualization, Methodology, Formal analysis, Investigation, Writing – original draft; Bilan Li: Methodology, Investigation, Validation, Writing – original draft. Funding acquisition; Ruru Zhao: Investigation, Resources, Data Curation, Writing – original draft; Huan Tong: Resources, Supervision, Writing – review & editing; Yinyan He: Formal analysis, Visualization, Writing – review & editing; Jing Ding: Conceptualization, Funding acquisition, Supervision, Project administration, Writing – review & editing. Guiqiang Du, Bilan Li, Yinyan He and Jing Ding have accessed and verified the underlying data. All authors have read and approved the final version of the manuscript and ensured that this is the case.

## References

1. Li Q-S, Zheng P-S. ESRRB Inhibits the TGFβ Signaling Pathway to Drive Cell Proliferation in Cervical Cancer. Cancer Res. 2023;83(18):3095–114.

2. Oh E-T, Kim HG, Kim CH, Lee J, Kim C, Lee J-S, et al. NQO1 regulates cell cycle progression at the G2/M phase. Theranostics. 2023;13(3):873–95.

3. Wang X, Jiang W, Du Y, Zhu D, Zhang J, Fang C, et al. Targeting feedback activation of signaling transduction pathways to overcome drug resistance in cancer. Drug Resist Updat. 2022;65:100884.

4. Javed Z, Muhammad Farooq H, Ullah M, Zaheer Iqbal M, Raza Q, Sadia H, et al. Wnt Signaling: A Potential Therapeutic Target in Head and Neck Squamous Cell Carcinoma. Asian Pac J Cancer Prev. 2019;20(4).

5. Takahashi-Yanaga F, Kahn M. Targeting Wnt signaling: can we safely eradicate cancer stem cells? Clin Cancer Res. 2010;16(12):3153–62.

6. Zhao H, Ming T, Tang S, Ren S, Yang H, Liu M, et al. Wnt signaling in colorectal cancer: pathogenic role and therapeutic target. Mol Cancer. 2022;21(1):144.

7. Wu X, Que H, Li Q, Wei X. Wnt/β-catenin mediated signaling pathways in cancer: recent advances, and applications in cancer therapy. Mol Cancer. 2025;24(1):171.

8. Zhou Y, Xu J, Luo H, Meng X, Chen M, Zhu D. Wnt signaling pathway in cancer immunotherapy. Cancer Lett. 2021;525:84–96.

9. Yu F, Yu C, Li F, Zuo Y, Wang Y, Yao L, et al. Wnt/β-catenin signaling in cancers and targeted therapies. Signal Transduct Target Ther. 2021;6(1):307.

10. Yadav UP, Singh T, Kumar P, Sharma P, Kaur H, Sharma S, et al. Metabolic Adaptations in Cancer Stem Cells. Front Oncol. 2020;10:1010.

11. Wang X, Chen Y, Wang X, Tian H, Wang Y, Jin J, et al. Stem Cell Factor SOX2 Confers Ferroptosis Resistance in Lung Cancer via Upregulation of SLC7A11. Cancer Res. 2021;81(20):5217–29.

12. Zeng Z, Fu M, Hu Y, Wei Y, Wei X, Luo M. Regulation and signaling pathways in cancer stem cells: implications for targeted therapy for cancer. Mol Cancer. 2023;22(1):172.

13. Pan G, Thomson JA. Nanog and transcriptional networks in embryonic stem cell pluripotency. Cell Res. 2007;17(1):42–9.

14. Belur Nagaraj A, Knarr M, Sekhar S, Connor RS, Joseph P, Kovalenko O, et al. The miR-181a-SFRP4 Axis Regulates Wnt Activation to Drive Stemness and Platinum Resistance in Ovarian Cancer. Cancer Res. 2021;81(8):2044–55.

15. Chen M-W, Yang S-T, Chien M-H, Hua K-T, Wu C-J, Hsiao SM, et al. The STAT3-miRNA-92-Wnt Signaling Pathway Regulates Spheroid Formation and Malignant Progression in Ovarian Cancer. Cancer Res. 2017;77(8):1955–67.

16. Li J, Yang S, Su N, Wang Y, Yu J, Qiu H, et al. Erratum to: Overexpression of long non-coding RNA HOTAIR leads to chemoresistance by activating the Wnt/β-catenin pathway in human ovarian cancer. Tumour Biol. 2015;36(11):9093–4.

17. Sankar S, Tanner JM, Bell R, Chaturvedi A, Randall RL, Beckerle MC, et al. A novel role for keratin 17 in coordinating oncogenic transformation and cellular adhesion in Ewing sarcoma. Mol Cell Biol. 2013;33(22):4448–60.

18. Luo Y, Pang B, Hao J, Li Q, Qiao P, Zhang C, et al. Keratin 17 covalently binds to alpha-enolase and exacerbates proliferation of keratinocytes in psoriasis. Int J Biol Sci. 2023;19(11):3395–411.

19. Baraks G, Tseng R, Pan C-H, Kasliwal S, Leiton CV, Shroyer KR, et al. Dissecting the Oncogenic Roles of Keratin 17 in the Hallmarks of Cancer. Cancer Res. 2022;82(7):1159–66.

20. Jang T-H, Huang W-C, Tung S-L, Lin S-C, Chen P-M, Cho C-Y, et al. MicroRNA-485-5p targets keratin 17 to regulate oral cancer stemness and chemoresistance via the integrin/FAK/Src/ERK/β-catenin pathway. J Biomed Sci. 2022;29(1):42.

21. Liang W, Liu H, Zeng Z, Liang Z, Xie H, Li W, et al. KRT17 Promotes T-lymphocyte Infiltration Through the YTHDF2-CXCL10 Axis in Colorectal Cancer. Cancer Immunol Res. 2023;11(7):875–94.

22. Dong Y, Wang B, Du M, Zhu B, Cui K, Li K, et al. Targeting Epsins to Inhibit Fibroblast Growth Factor Signaling While Potentiating Transforming Growth Factor-β Signaling Constrains Endothelial-to-Mesenchymal Transition in Atherosclerosis. Circulation. 2023;147(8):669–85.

23. Brophy ML, Dong Y, Tao H, Yancey PG, Song K, Zhang K, et al. Myeloid-Specific Deletion of Epsins 1 and 2 Reduces Atherosclerosis by Preventing LRP-1 Downregulation. Circ Res. 2019;124(4).

24. Song K, Cai X, Dong Y, Wu H, Wei Y, Shankavaram UT, et al. Epsins 1 and 2 promote NEMO linear ubiquitination via LUBAC to drive breast cancer development. J Clin Invest. 2021;131(1).

25. Li W, Chen C, Zheng H, Lin Y, An M, Liu D, et al. UBE2C-induced crosstalk between mono- and polyubiquitination of SNAT2 promotes lymphatic metastasis in bladder cancer. J Clin Invest. 2024;134(13).

26. Zhang P, Gao K, Zhang L, Sun H, Zhao X, Liu Y, et al. CRL2-KLHDC3 E3 ubiquitin ligase complex suppresses ferroptosis through promoting p14ARF degradation. Cell Death Differ. 2021;29(4):758–71.

27. Wang H, Peng J, Li H, Lan Y, Guo J, Qiu Q, et al. E3 Ubiquitin Ligases: Structures, Biological Functions, Diseases, and Therapy. MedComm (2020). 2025;6(12):e70528.

28. Shang F, Deng G, Liu Q, Guo W, Haas AL, Crosas B, et al. Lys6-modified ubiquitin inhibits ubiquitin-dependent protein degradation. J Biol Chem. 2005;280(21):20365–74.

29. Shaeffer JR, Cohen RE. Differential effects of ubiquitin aldehyde on ubiquitin and ATP-dependent protein degradation. Biochemistry. 1996;35(33):10886–93.

30. Kirfel J, Magin TM, Reichelt J. Keratins: a Structural Scaffold with Emerging Functions. Cell Mol Life Sci. 2003;60:56–71.

31. Nair RR, Hsu J, Jacob JT, Pineda CM, Hobbs RP, Coulombe PA. A role for keratin 17 during DNA damage response and tumor initiation. Proc Natl Acad Sci U S A. 2021;118(13).

32. Wickramarachchi DC, Theofilopoulos AN, Kono DH. Immune pathology associated with altered actin cytoskeleton regulation. Autoimmunity. 2010;43(1):64–75.

33. Naydenov NG, Koblinski JE, Ivanov AI. Anillin is an emerging regulator of tumorigenesis, acting as a cortical cytoskeletal scaffold and a nuclear modulator of cancer cell differentiation. Cell Mol Life Sci. 2020;78(2):621–33.

34. Song P, Gao Z, Bao Y, Chen L, Huang Y, Liu Y, et al. Wnt/β-catenin signaling pathway in carcinogenesis and cancer therapy. J Hematol Oncol. 2024;17(1):46.

35. Clevers H, Nusse R. Wnt/β-catenin signaling and disease. Cell. 2012;149(6):1192–205.

36. Liu J, Smith S, Wang C. Photothermal Attenuation of Cancer Cell Stemness, Chemoresistance, and Migration Using CD44-Targeted MoS₂ Nanosheets. Nano Lett. 2023;23(5):1989–1999.

37. Fei Y, Cao D, Li Y, Wang Z, Dong R, Zhu M, Gao P, Wang X, Cai J, Zuo X. Circ_0008315 Promotes Tumorigenesis and Cisplatin Resistance and Acts as a Nanotherapeutic Target in Gastric Cancer. J Nanobiotechnol. 2024;22:519.

