## Supplementary table1-2 and Supplementary Figure 1-3 for "KRT17 stabilizes EPN1 via inhibiting SMURF1-mediated ubiquitination to modulate Wnt/β-catenin signaling output and stem-like traits in ovarian cancer"

### Supplementary data

**Table S1 The target sequences of shRNA in the genes FAM117B, KAZALD1, KRT17, LPHN2, and ZNF608.**

| Genes | Target sequence 1 | Target sequence 1 | Target sequence 3 |
|---|---|---|---|
| ZNF608 | AAGGAGGAAACTAAACAGAAA | / | / |
| KRT17 | ACAGCCAGTACTACAGGACAA | GCGTGACCAGTATGAGAAGAT | CACCTGACTCAGTACAAGAAA |
| KAZALD1 | GAGGAAGGATGGCTTGGACAT | / | / |
| FAM117B | AGCAGTTGCAGAGAAGTAAAC | / | / |
| ADGRL2 | ACAAAGAAAGAACGAGGAATA | / | / |

**Table S2 Primers for real-time PCR.**

| Genes | Forward(5′–3′) | Reverse(5′–3′) |
|---|---|---|
| GAPDH | TGACTTCAACAGCGACACCCA | CACCCTGTTGCTGTAGCCAAA |
| KRT17 | CCGTCTGGCTGCTGATGACT | CTCCTCCTCGTGGTTCTTCTTCA |
| EPN1 | CAAGAACTGGCGTCACGTTTA | CTGCTTAGCTTTCTCACGCA |

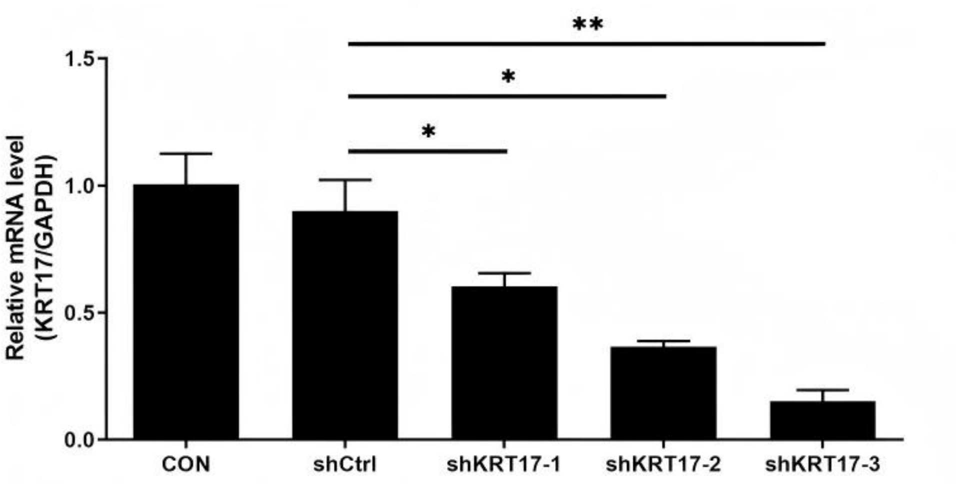


**Supplementary Figure 1 Screening for effective interference targets for KRT17 by qRT-PCR.**

The expression levels of KRT17 RNA in SK-OV-3 cells transfected with shCtrl and shKRT17 (1 - 3) plasmids were measured using qRT - PCR.*P < 0.05; **P < 0.01.

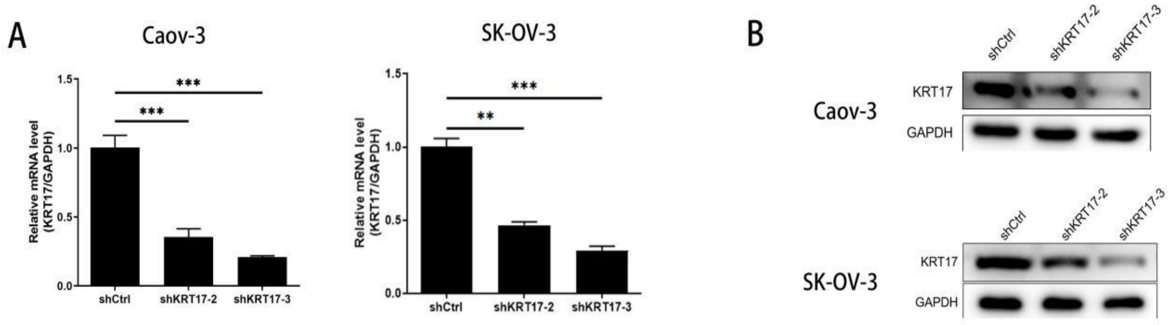


**Supplementary Figure 2 The knockdown efficiency of KRT17 in OC cells at the mRNA and protein levels.**

**A - B:** The expression levels of KRT17 RNA and protein in Caov - 3 and SK - OV - 3 cells stably transfected with shCtrl and shKRT17 (1 - 2) plasmids were measured using qRT - PCR (**A**) and western blotting (**B**). **P < 0.01; ***P < 0.001;

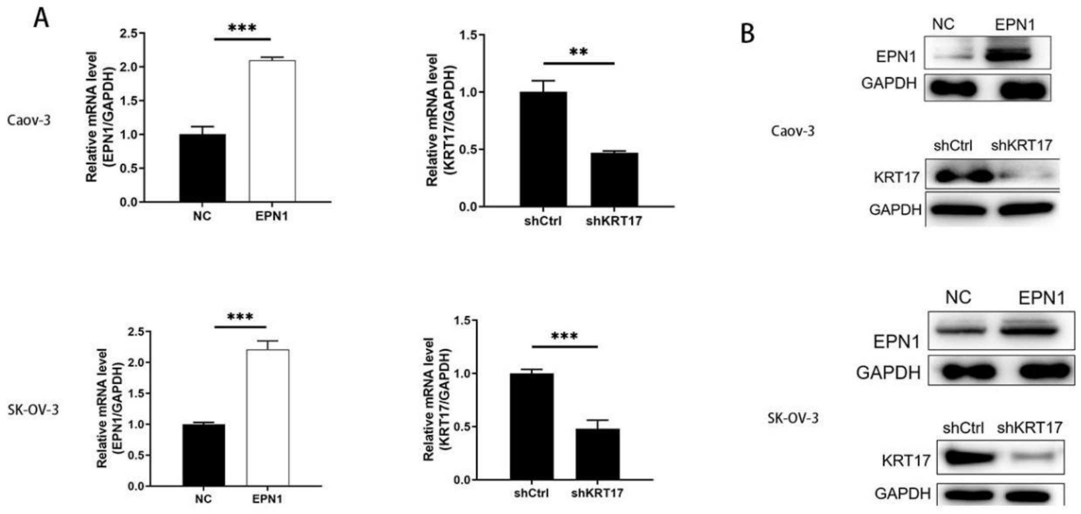


**Supplementary Figure 3 Transfection efficiencies of KRT17 and EPN1 in OC cells at the mRNA and protein levels.**

**A - B:** Validation of the expression levels of target gene RNA (**A**) and protein (**B**) in Caov - 3 and SK - OV - 3 cells stably expressing NC or EPN1, as well as shCtrl or shKRT17, was performed by qRT - PCR and immunoblotting.
